# Mosaic evolution, preadaptation, and the evolution of evolvability in apes

**DOI:** 10.1101/622134

**Authors:** Caroline Parins-Fukuchi

## Abstract

A major goal in post-synthesis evolutionary biology has been to better understand how complex interactions between traits drive movement along and facilitate the formation of distinct evolutionary pathways. I present analyses of a character matrix sampled across the haplorrhine skeleton that revealed several modules of characters displaying distinct patterns in macroevolutionary disparity. Comparison of these patterns to those in neurological development showed that early ape evolution was characterized by an intense regime of evolutionary and developmental flexibility. Shifting and reduced constraint in apes was met with episodic bursts in phenotypic innovation that built a wide array of functional diversity over a foundation of shared developmental and anatomical structure. Shifts in modularity drove dramatic evolutionary changes across the ape body plan in two distinct ways: 1) an episode of relaxed integration early in hominoid evolution coincided with bursts in evolutionary rate across multiple character suites; 2) the formation of two new trait modules along the branch leading to chimps and humans preceded rapid and dramatic evolutionary shifts in the carpus and pelvis. Changes to the structure of evolutionary mosaicism may correspond to enhanced evolvability that has a ‘preadaptive’ effect by catalyzing later episodes of dramatic morphological remodeling.

## Introduction

The formation of new species and phenotypes is often hypothesized to be limited by the availability of unoccupied ‘adaptive zones’, classically represented as peaks on a fitness landscape (Wright 1932, Simpson 1944). Frequently referred to as ‘ecological opportunity’ (Losos 2010), the opening of new adaptive zones through shifts in biotic interactions and the environment is often invoked to explain large evolutionary radiations (Rainey and Travisano 1998, Yoder et al., 2010, Wagner and Harmon et al 2012, but see Slater 2015). The role of ecological processes in generating morphological novelty can be contrasted with an increasing focus on ‘constructional’ factors that include functional interactions and architectural constraints (Seilacher 1970, 1991). While character evolution may sometimes reflect simple optimization processes when natural selection operates independently on separate traits, the evolutionary pathways realized in nature are also constrained by the complex interactions between functionally dependent traits and the limitations of an organism’s developmental architecture. When placed into a broader evolutionary literature, these limiting factors might be coarsely partitioned into developmental (Gould and Lewontin 1979, Alberch 1980, Olson 2012) and functional (Charlesworth et al., 1982, Maynard-Smith et al., 1985, Arnold 1992) constraints.

Developmental and functional constraints limit the capacity for a clade to evolve novel phenotypes. Together, these factors might be said to determine the level of ‘constructional opportunity’ available at a given time. Contrasted to ecological opportunity, constructional opportunity reflects how the structure of developmental and functional interactions between characters limits the ‘evolvability’ of a clade—its capacity to evolve novel phenotypes. While ecological opportunity is often evoked as a limiting factor on a clade’s phenotypic diversification, constructional opportunity, as measured by the overall level of constraint and internal integration between phenotypes, may also limit the range of evolutionary modifications available to a population faced with ecological pressure.

Evolutionary constraints are shaped by shared developmental pathways, multivariate selection, and underlying genetic processes such as epistasis and pleiotropy. Such interactions between traits and genes often create a common tendency to drive the formation of modules of characters that are internally integrated in their evolutionary trajectories. Such integration is typically identified by calculating numerical correlations in their variance (Olson and Miller 1958). Integration between traits constrains the combination of phenotypes accessible to each. For example, for two traits that are strongly positively integrated, it is unlikely to observe an individual that possesses phenotypes for each trait that lie at opposing ends of the spectrum of variation. As long as this correlation exists, the evolution of the organism is constrained in the sense that only certain combinations of phenotypes are accessible. At the population level, integration can be caused by multivariate selection or physical/constructional linkages such as pleiotropy or allometry (Cheverud 1984). However, it is generally not possible to distinguish between these in macroevolutionary study systems.

Phenotypic integration generally occurs in a modular fashion (Wagner et al. 2007). Instead of covarying uniformly across the entire organism, integration often manifests as a patchwork of modules of internally integrated traits that are free to evolve largely independently (Goswami et al. 2014, 2015). Such modular structures lead to the pattern of mosaic evolution when separate modules evolve at different rates as lineages diverge (Felice and Goswami 2017, Parins-Fukuchi, in press). When analyzed phylogenetically, these patterns result in ‘mosaic disparity’—a scenario where individual modules of traits each display a unique pattern in phylogenetic disparity across lineages (Fig. 1).

**Figure 1.**
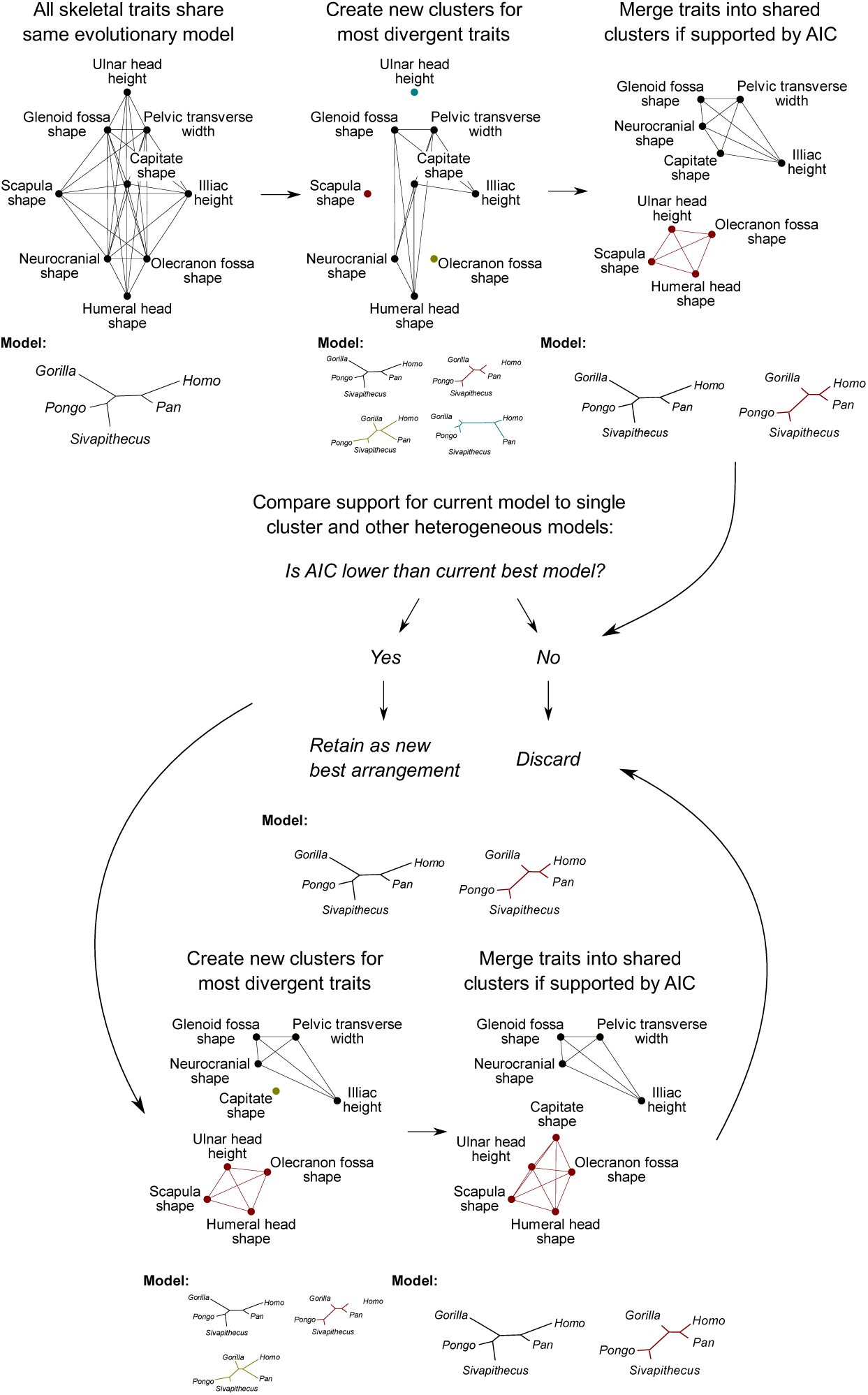
The mosaic inference procedure starts by modelling all characters under a unified phylogenetic model, in which branch lengths represent morphological disparity. The algorithm then splits traits with divergent patterns in disparity into their own suites and then greedily merges these suite fragments until the AIC score ceases to improve. The splitting and merging steps are then alternated until the AIC scores of the final merged models cease to improve. The best clustering achieved from this procedure represents the best-supported heterogeneous model of character integration.

The related concepts of mosaicism, constraint, and integration can help to explain the emergence of phenotypic novelty across clades. For example, when ecological opportunity is abundant, evolutionary radiations can be driven primarily by the release of constraints (Wagner et al. 2003). In addition, bursts in evolutionary rate can also coincide with the break-up of existing modules and the subsequent formation of a new modular structure (Wagner 2018). Characterizing the role of integration and mosaic evolution in shaping diversity across clades will ultimately contribute to a deeper understanding of phenotypic evolution at the constructional level. This may improve our understanding of evolutionary processes more generally by revealing the extent to which fluctuations in constructional opportunity contribute to macroevolutionary patterns alongside ecological opportunity.

In this study, I sought to determine the extent to which the evolutionary trajectories blazed by ape species were shaped by early shifts in the structure of integration and the timing of development. To address these questions, I reconstructed the mosaic macroevolutionary patterns in morphological, neurological, and developmental phenotypes across living and fossil great apes throughout the Miocene. To help place these empirical results in a theoretical context, I also present a set of simple simulations to generate a theoretical expectation for the patterns in disparity expected when Markovian diffusion is constrained by trait integration imposed by both functional and structural/developmental interactions. Taken together, these analyses suggest the capability for shifting patterns in constraint imposed by integration to create constructional opportunities that generate repeated and dramatic episodes in morphological innovation.

Apes have been extensively studied in the context of both developmental constraint and environmental adaptation. Young et al. (2010), found that apes display weak integration between the fore- and hindlimb in comparison to cercopithecoids. Reduced integration can confer greater functional flexibility, and so may have facilitated the proliferation of the diverse locomotor function across living and fossil apes. Changes in developmental timing have also been important in ape evolution. Developmental heterochrony appears to have driven the emergence of many key aspects of hominin morphology in the cranium and post-cranium (Gould 1977, Berge 1998). This long-standing interest in the developmental basis of ape evolution has not excluded environmental hypotheses. Environmental fluctuations throughout the Miocene and Pliocene have also frequently formed the basis for adaptive hypotheses across apes (Andrews 1992, Ruff 1994, Michel et al., 2014). The body plans of hominins, and undoubtedly apes in general, have been shaped through modifications to separate suites of traits occurring at different times (Holloway 1973, McHenry 1975). These diverse threads suggest that a multiplicity of causes, perhaps including both ecological opportunity and constructional factors, has shaped the evolution of hominoid body plans. As a result, apes are an excellent exemplar taxon in which to pluralistically examine the mosaic patterns in constraint and innovation that shape body plans in living and fossil vertebrates.

## Methods

### Morphological data

I gathered a dataset of 149 quantitative morphological traits from the literature (Lewton 2010, Worthington 2012) spanning the cranium, forelimb, and pelvis. These traits were sampled across 10 extant taxa. I also collected fossil cranial and forelimb traits from Worthington (2012), retaining all taxa with at least 25% matrix occupancy. Taxa were all studied at the generic level, with each extant genus represented by a single exemplar species. Each morphological trait represented either a dimensionless ratio or geometric mean, depending on how the author of each dataset handled intraspecific variation and body mass correction. I scaled the traits to display an empirical variance of 1 across all taxa. This simplified the analyses by reducing the complexity of the dataset, while retaining the same comparative information. Since the original traits were dimensionless, both the unscaled and the scaled datasets would facilitate examination of *relative,* rather than absolute, evolutionary rates, but the scaled traits improve Markov-chain Monte Carlo (MCMC) mixing and simplify the identification of mosaic suites.

### Neurological data

To directly study the evolutionary patterns in related developmental and phenotypic traits, I also gathered neurological data from the literature (Capellini et al., 2010, Boddy et al., 2012). For neurological phenotypes, I collected encephalization quotients (EQ) estimated in 76 primate species by Boddy et al. (2012). EQ is calculated by fitting a nonlinear model to brain size and body size and taking the deviations (residuals) from the best fit curve and so measures the enlargement of brain size while controlling for allometry with body size. I compared EQ patterns to those in a postpartum encephalization development (ED) metric calculated as:

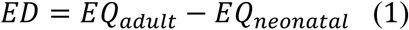

ED is therefore intended to reflect the amount of brain tissue grown by members of each species following birth relative to their body size. The ED metric was scaled to unit variance across all sampled taxa. Neonatal brain and body mass data representing 24 primate species were acquired from Capellini et al. (2010).

### Fossil placement and divergence time estimation

Since the evolutionary relationships between the extant taxa represented in the morphological dataset have all been extensively studied and are well-resolved, I used these as a fixed scaffolding to place the fossil taxa. Fossil placements were inferred from the continuous trait dataset using the cophymaru program (Parins-Fukuchi 2018). I used the ‘binary weights’ procedure to filter out reliable traits from ones likely to mislead by using the extant tree as a fixed point of reference. Markov-chain Monte Carlo (MCMC) simulations were run for 1,000,000 generations and checked manually for convergence. The posterior tree sample was summarized as the maximum clade credibility tree using sumtrees.py (Sukumaran and Holder 2010).

For downstream comparative analyses, I also estimated divergence times on the full 19-taxon tree. I downloaded cytochrome B sequences from Genbank representing each of the 10 extant taxa. Molecular dating was performed using Beast version 2.4.8 (Bouckaert et al. 2014) using the fossilized birth-death (FBD) prior. The topology was fixed to the summary tree generated during the fossil placement step. Temporal occurrence ranges for the fossil taxa were assembled from the literature. These fossil and extant occurrence ranges were used in the dating analysis to infer the diversification and sampling rate parameters used in the FBD prior.

### Measuring integration throughout the hominoid radiation

As an evaluation of the overall strength of integration in the skeletal traits across the haplorrhine phylogeny, I estimated ancestral states for each character under Brownian motion along a dated tree of nine extant haplorrhine taxa (*Pan paniscus* was removed because it was missing pelvic measurements). I then estimated the strength with which the reconstructed phenotypes at each internal node were correlated with the values at each descendant node or tip. The correlations indicated how strongly the reconstructed evolutionary changes in skeletal traits were integrated along each branch in the tree. Stronger correlations corresponded to tighter overall integration. The Python script that I developed for the reconstruction of ancestral states and calculation of character correlations is available in the data supplement.

### Evolutionary rates across mosaic morphological suites

I used the *greedo* program to recover mosaic patterns in disparity from the continuous trait dataset (Parins-Fukuchi 2019). *Greedo* is a phylogenetic clustering/mixture model estimation approach that uses the Akaike information criterion (AIC) to iteratively merge and split clusters of traits to find a best-supported set of character suites based on shared patterns in disparity (i.e., where integrated traits within the same module are similarly disparate across taxa), where each is represented by a tree with branch lengths (Fig. 1). Traits that display the highest improvement in log-likelihood when assigned to a separate cluster are prioritized during the splitting steps to first identify likely sources of heterogeneity in the dataset. During the merging step, only clusters that show an improved AIC score when placed under the same model are joined. This process recovers an estimate of the number of suites, the membership of each trait, and a tree with branch lengths scaled to units of disparity for each suite. Fossil taxa were not included in the mosaic analyses, because they were too fragmentary in their sampling to inform disparity patterns when the traits were split into suites. I performed many runs of the *greedo* procedure to avoid presenting results obtained from a suboptimal peak on the likelihood surface. Since several of the top clusterings yielded close AIC support, I performed an additional model-averaging step using Akaike weights to summarize the results as a graph where each trait occupies a node where each link is weighted to reflect the total weighted AIC support across all the models. Comparative analyses were performed using a single summary clustering that resulted from application of the Markov clustering (MCL) algorithm to the support network (Dongen 2000).

To examine the anatomical composition of the mosaic suites of characters, I performed a statistical enrichment procedure to identify skeletal regions that were over and under-represented in each suite relative to the proportions for each present in the entire dataset. The proportion of the 149-trait dataset represented by each skeletal region was used to generate a set of ‘expected’ values. I then compared these to the observed proportions within each suite to examine trends in the deviations of the proportions occupied by each region. I did not assess the statistical significance of these deviations, because 1) my aim was to examine the general patterns in the composition of each character suite rather than to determine absolute significance, and 2) the small sample size of the dataset generated expected chi-square test statistic values that were nearly entirely <5, making the test inappropriate. To examine the relative contributions of separate anatomical regions in shaping the macroevolutionary patterns reconstructed in each suite, I also performed a principal component analysis (PCA) on the traits contained within each suite. I then transformed the loadings to calculate the total variance contributed by each trait across all axes of the PCA, summing them to produce the contribution of each anatomical region.

I transformed the branch lengths for each mosaic suite, which were scaled to reflect disparity (total variance, *v*, accumulated over time *t*), to rates (*v* per unit *t*) using the results from the divergence time analysis. I also calculated evolutionary rates averaged over the entire morphological dataset while including the fossil taxa to coarsely reconstruct evolutionary tempo throughout the Miocene. Although this step missed valuable information recovered by the mosaic analyses, including the fossil taxa in this manner enabled finer resolution into the coarse patterns in evolutionary tempo throughout the Miocene.

### Evolutionary rates in neurological data

I estimated macroevolutionary rates across primates in ED and EQ using BAMM version 2.5.0 (Rabosky 2014). Since the trees that I constructed for the morphological analysis contained only a fraction of the taxa present in the neurological datasets, I used the dated primate supertree packaged in BAMM that was originally sourced from Mos and Mooers (2006) and pruned the tips to match the taxa present in each dataset. MCMC simulations were run until the estimated sample size (ESS) well exceeded 200 for each parameter. Results were presented by plotting the mean rate estimated along each branch and the maximum *a posteriori* configuration of rate shifts using BAMMtools version 2.5.0 in R (Rabosky et al., 2014).

### Lineage diversification rate analyses

I performed analyses of origination and extinction rates in hominoid and cercopithecoid fossil records to examine the correspondence between lineage diversification and the patterns in disparification emphasized in the analyses of morphological rates and mosaicism. Fossil occurrence data spanning primates were acquired from the Paleobiology Database (paleobiodb.org) on July 31, 2019. This dataset was partitioned into two subsets: one encompassing Hominoidea and the other representing Cercopithecoidea. Rates of origination and extinction were inferred from each subset separately using PyRate (Silvestero et al. 2014). The inferential model was constrained to contain uniform origination and extinction rates across time, thus ignoring the possibility of diversification shifts within each clade. Preservation was constrained to being time-homogeneous and was assumed to be uniform within each clade. While ignoring potential information regarding diversification rate heterogeneity through time, this simplified model matches classic paleobiological work that assumes simple birth-death-sampling models and was therefore appropriate for the comparison of average diversification rates in sister clades while avoiding possible model overfitting that may obfuscate the straightforward test sought here.

### Simulated Markovian diffusion

To examine the expected effect of constructional opportunity stemming from both developmental and functional constraint on phenotypic evolution, I designed a set of evolutionary simulations based on a simple Markovian diffusion. In this system, quantitative traits belonging to a single population of organisms evolve stochastically along a fitness landscape. Each generation, values for each trait are proposed. Values that increase the overall fitness of the population are accepted. To mimic the effect of drift, values that decrease fitness are also accepted with some (user-specified) probability. Constraint stemming from developmental integration caused by pleiotropy can be mimicked by proposing values for traits that covary, while functional constraints are modeled by evolving traits along a shared, multivariate adaptive landscape, modelled here using a mixture of multivariate Gaussian distributions. In a completely unconstrained system, each trait evolves according to its own univariate landscape, with its value drawn independently of all others in each generation. Performing simulations in this way facilitated an illustration of the potential for functional and developmental integration to constrain evolutionary rates in an adaptive landscape. Although distinguishing between these modes of integration is generally difficult or impossible in paleontological study systems, the simulations performed here provide insight by testing 1) the potential for varying systems of integration to catalyze or constrain adaptive change and 2) the extent to which functional and developmental integration generate overlapping and distinct patterns in evolutionary rate and disparity. When placed in a macroevolutionary context, the simulations here should be thought of as a demonstration of the possible microevolutionary processes that may have driven the higher-level patterns in evolutionary rate occurring along a single phylogenetic branch in the mosaic analyses. The script used to generate these simulations is available in the data supplement.

## Results and Discussion

### Strength of integration throughout the haplorrhine radiation

Calculating character correlations from the skeletal data along each branch of the extant haplorrhine phylogeny revealed a single episode of reduced integration at the root of Hominoidea, where the correlation decreased from 0.12 at the ancestor of catarrhines, to −0.04 at the root of hominoids (Fig. 2). This suggests that the reduction in integration across apes between the fore- and hindlimbs revealed by Young and colleagues (2009) extended across the skeleton along the branch leading to the stem hominoids. Such episodic relaxed integration was not detected in the other haplorrine lineages, with cercopithecoids (0.2-0.33 correlation) and platyrrhines (0.16-0.31 correlation) displaying higher integration than hominoids (−0.04-0.25 correlation) overall. However, it is possible that more complete sampling of taxa and characters across both of these clades may reveal changes in modularity and integration not detected here.

**Figure 2.**
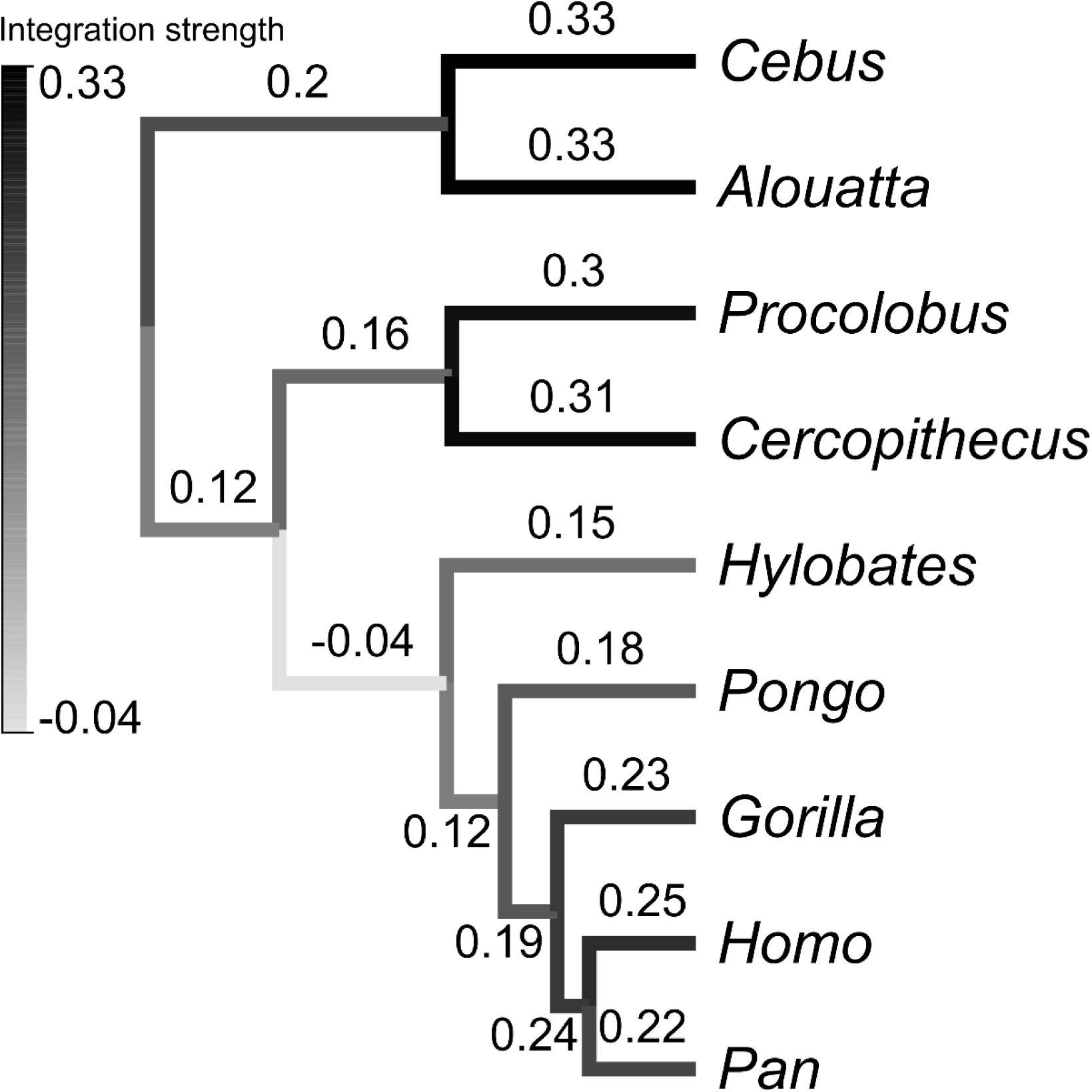
Strength of integration across haplorrhine evolution. Branch labels correspond to the strength and polarity of correlation in the reconstructed changes along each branch. Higher correlations in either direction (closer to either 1 or −1) indicate stronger evolutionary integration, while lower correlations (closer to 0) indicate weaker integration.

### Composition of mosaic suites

The analysis of mosaic disparity recovered five suites of traits (Fig 3b). The cranium was the largest contributor of both traits and variance across all suites except for C4, which was represented by the pelvis alone (Fig. 3b and 3c). This reflects the higher sampling of cranial traits in the dataset. While the cranium has previously been shown to display substantial modularity (Felice and Goswami 2017), the results here suggest that individual cranial modules may result from the formation of broader evolutionary complexes shared with the post-cranium. The suites were distinct in their composition across postcranial anatomy: C0 was represented strongly by the wrist, ulna, and humerus; C1 was represented postcranially by the scapula; C2 by carpal and pelvic traits. The cranium was represented even more strongly than expected in C3 and contributed nearly half of the variance (Fig. 3b).

**Figure 3.**
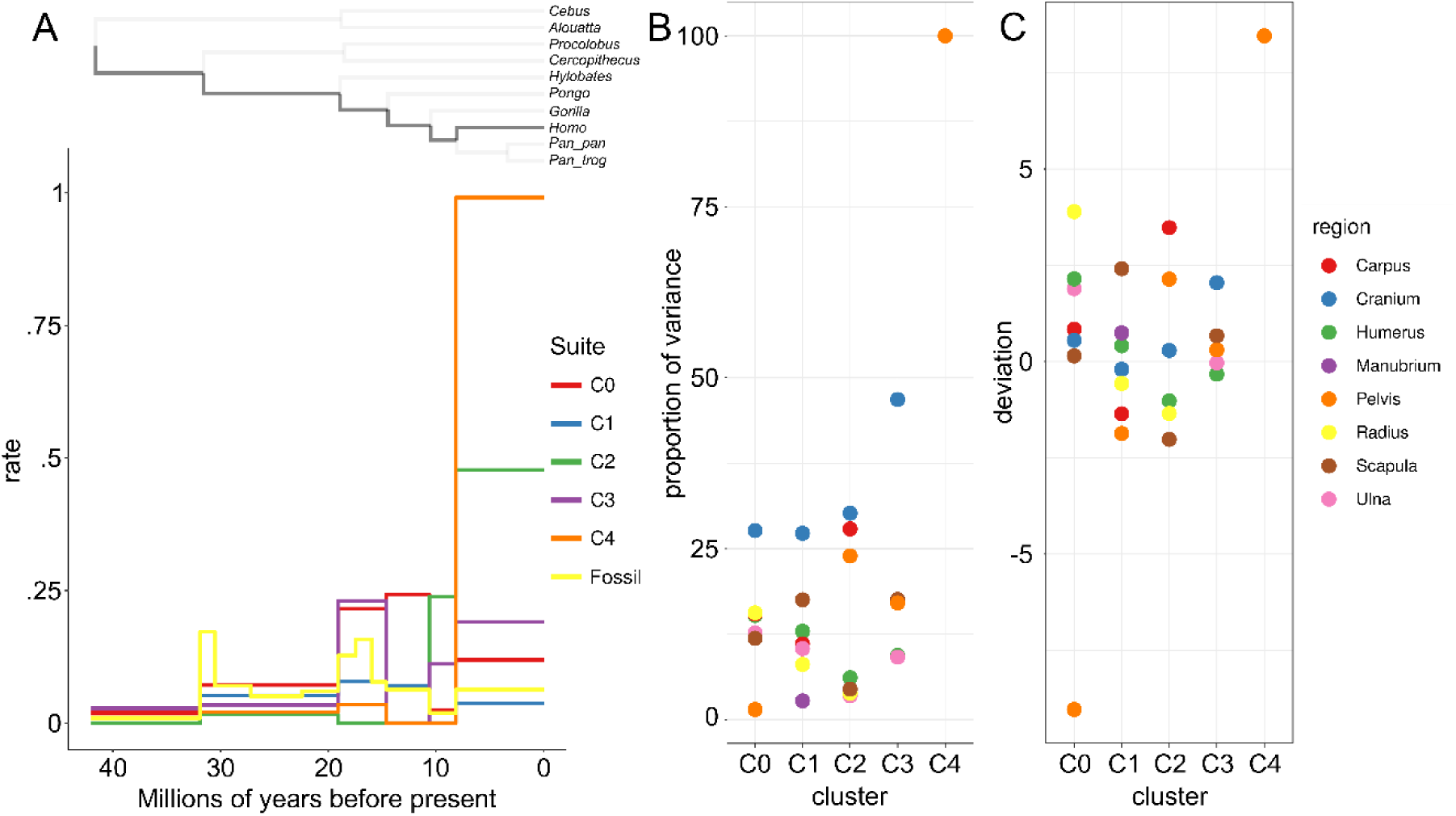
A) Mosaic evolutionary rates calculated while walking back from *Homo* to the most recent common ancestor of New World monkeys, Old World Monkeys, and apes. B) Variance contributed to each suite by each skeletal region, estimated from a principal component analysis. C) Skeletal regions over-represented in each mosaic evolutionary suite. Values above zero correspond to regions that occupy a higher proportion of their clusters than expected.

### Rates of morphological evolution

The phylogenetic rate calculations for each suite revealed a strong pattern of evolutionary mosaicism (Fig. 3a). Three of the suites, C0, C1, and C3, displayed the highest rates after the divergence of the great apes from hylobatids, but before the divergence of gorillas from humans and chimpanzees. The two suites that did not experience the shared great ape rate increase, C2 and C4, experienced large bursts in *Homo*. This finding is consistent with general knowledge of human evolution, as both suites are dominated by cranial traits and pelvic traits related to birthing and locomotor function (Table S1).

**Table 1.** Lineage diversification and preservation rate parameters estimated in apes (Hominoidea) and Old World monkeys (Cercopithecoids). Parameters were summarized as the mean values from four million Markov-Chain Monte Carlo generations after discarding 20% as burn-in.

| Clade | Origination | Extinction | Net diversification | Preservation |
| --- | --- | --- | --- | --- |
| Hominoidea | 0.395 | 0.338 | 0.057 | 3.121 |
| Cercopithecoidea | 0.251 | 0.128 | 0.123 | 2.551 |

The rate analyses (Fig. 3a) show that humans have experienced major macroevolutionary bursts. However, the unique characteristics of the human body plan are built over an anatomical structure shared by all great apes that was shaped in the Miocene during similarly dramatic episodes. For example, although suite C0 increased in evolutionary rate along the human branch, its evolution was shaped by a sustained period of elevated rate between 20 and 10 million years ago. This suggests that much of the ‘groundwork’ underlying the derivation of humans’ unique features may predate the divergence between our lineage and chimpanzees’. The mosaic analysis also demonstrates that substantial visible phenotypic novelty can result from the evolutionary remodeling of a small subset of the anatomy. The most elevated evolutionary rates in humans were observed in suites C2, C3, and C4, which cumulatively comprise fewer than one-third of the traits sampled in the matrix (Table S1). C4, the smallest suite (10 traits) displayed by far the largest increase in rate along the *Homo* lineage. Perhaps notably, C4 also was the most static character suite throughout the earlier stages in the hominoid radiation. The mosaic pattern recovered here shows that the uniqueness of the human body plan was shaped by dramatic evolutionary shifts in only a small number of traits.

While the large amount of missing data among the fossil taxa made an additional mosaic analysis infeasible, their inclusion revealed an otherwise hidden shift in evolutionary rate that occurred at the root of Hominoidea (approximately 30 million years ago). The fossil data also recapitulated the burst in evolutionary rate during the mid-Miocene that was displayed by C0 and C1 in the mosaic analysis (Fig. 3 and 4). When averaging over all the traits and including fossil taxa, these two episodes are the most dramatic macroevolutionary events when tracing the evolutionary lineage leading to *Homo*, suggesting the importance of early shifts in the ape body plan in shaping the functional morphologies of living taxa.

**Figure 4.**
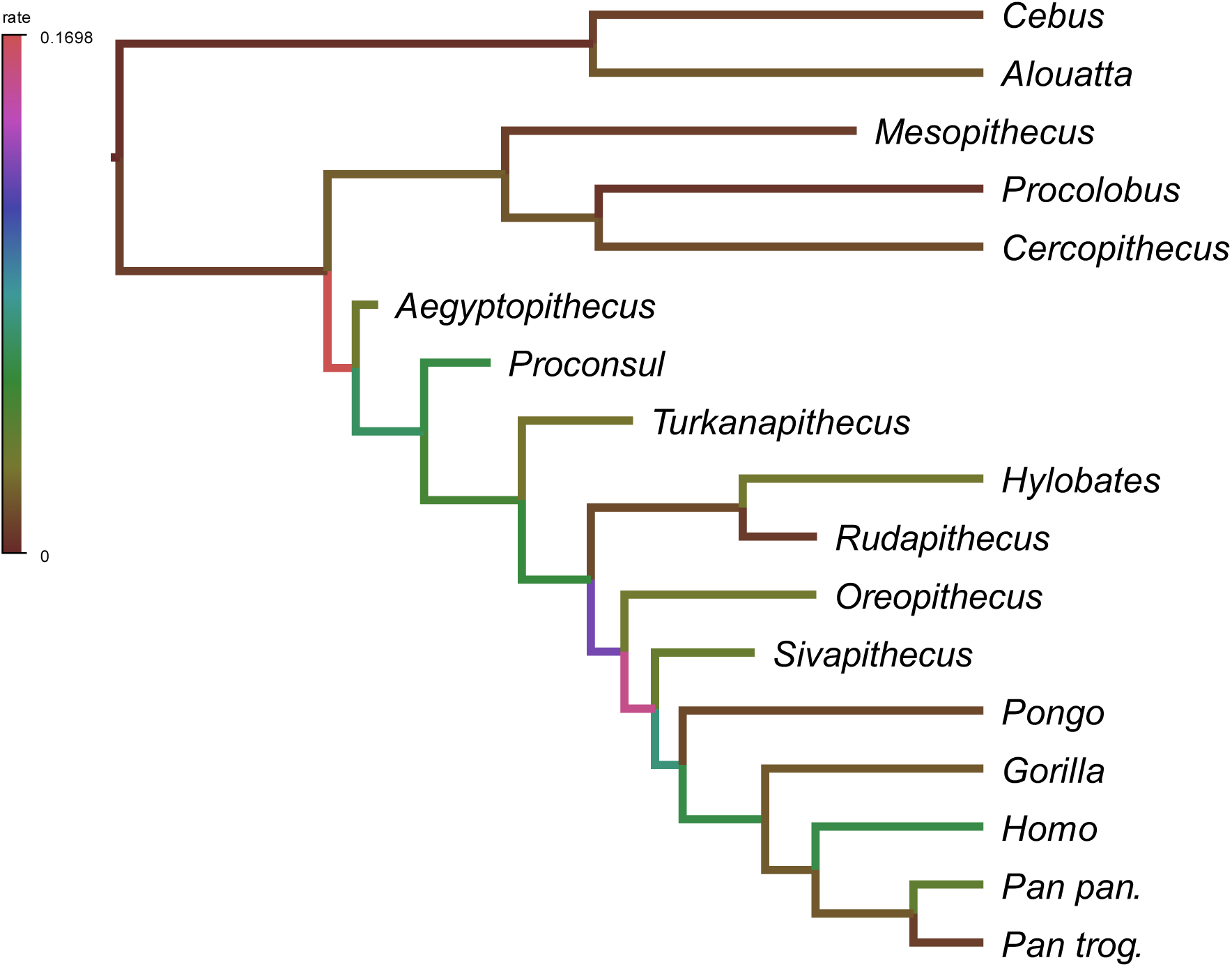
Branch colors correspond to rates of morphological evolution across all characters along individual branches. Phylogenetic positions of fossil genera reflect placements on extant reference tree.

Several of the major rate shifts that occurred throughout the great ape clade corresponded to increased evolvability conferred by relaxed integration among early apes. However, comparison of the human-specific evolutionary rates in cranial, pelvic, and neurological traits to the phylogenetic character correlations in Figure 2 shows that the elevated morphological rates in humans were immediately preceded by an *increase* in the strength of integration. This illustrates an alternative means through which shifts in systems of modularity promote rapid and dramatic evolutionary remodeling of morphology. The suites that showed the most dramatic rate increases in humans, C2 and C4, were evolutionarily inactive prior to the most recent common ancestor (MRCA) of chimps and humans. This suggests that, while early ape evolution was characterized by relaxed integration leading to rapid evolutionary change in existing modules, human morphological evolution was shaped by the formation of two small modules of characters from the pelvis and carpus that underwent an initial rate increase following their formation along the branch leading to chimps and humans and then a more dramatic rate increase in hominins. Although the overall structure of integration was well-defined along the chimp-human MRCA branch, the formation of *new*, smaller modules contributed to dramatic changes across the body plan in humans. The occurrence of these initial shifts in modularity and rate along the chimp-human MRCA shows that chimpanzees share the modified structure of integration that facilitated the evolution of key human traits. This result is surprising given the typical assumption that chimpanzees are morphologically more similar to other great apes than are humans.

### Rates of neurological evolution

The great apes differentiated rapidly from other catarrhines in both EQ and ED. *Homo* displays the highest rate of both EQ and ED evolution. However, the human rate shift in both traits occurred as part of an older trend of rapid neurological evolution in African apes. For EQ, humans experienced a substantial increase in evolutionary rate relative to the rest of the African apes, despite the statistical evidence for a shared rate shift at the root of the clade (Fig. 5b). The shared macroevolutionary regime shared by all African apes is most apparent in encephalization development (Fig. 5a), where the estimated evolutionary rate in *Homo* increased only slightly after splitting from *Pan* (Fig. 5c). It appears that the increased encephalization developed throughout the post-natal period in humans reflects a general trend among great apes in the evolutionary plasticity of neurological development. As a result, the ability to develop a relatively large mass of brain tissue does not itself appear to be a strong limiting factor in the evolution of large encephalization in humans. Instead, the morphological analyses suggest that pelvic traits (C4 in Fig. 1) demanded a more dramatic alteration in macroevolutionary regime, requiring both the formation of a new suite of characters and massive burst in rate.

**Figure 5.**
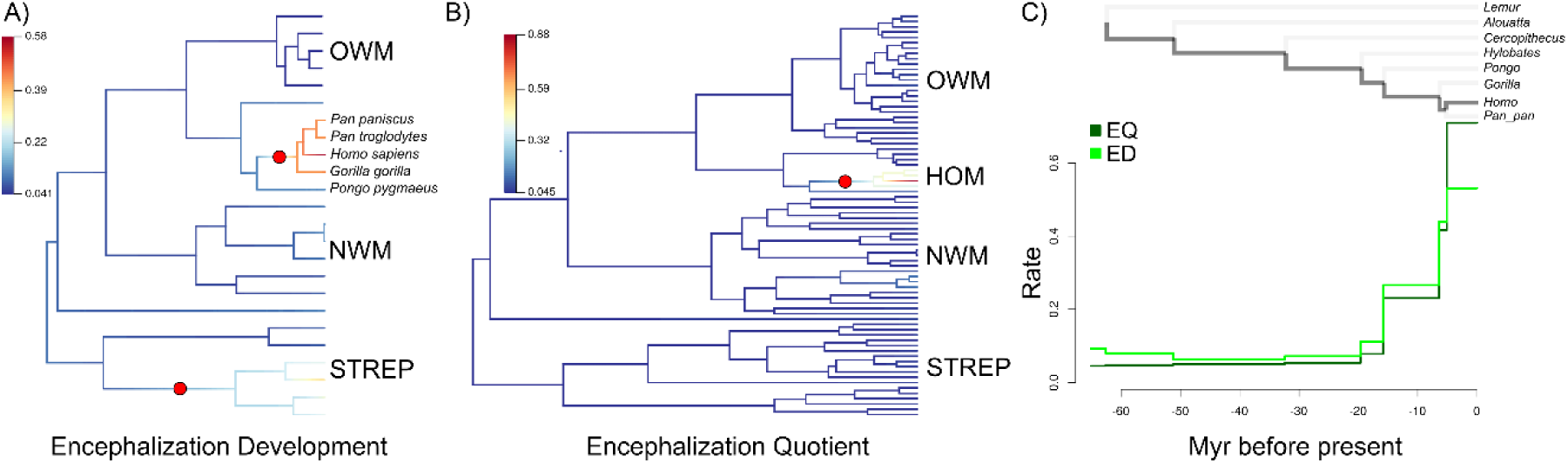
Rate shifts in A) encephalization development, and B) encephalization quotient. C) Branch-specific evolutionary rates when tracing the lineage from *Homo* to the root.

### Structural and ecological opportunity in ape evolution

The elevated evolutionary rates experienced by apes early in their divergence correspond to a general relaxation of constraint in early Miocene stem and ancestral hominoids. The reduction in integration across the skeleton and burst in the evolutionary rate of neurological development that occur at the base of the hominoid clade correspond in timing to the elevated rates in morphological evolution observed in most of the skeletal characters examined here (Fig. 3 and 4). Freed from previous functional and developmental limitations, ape body plans would have rapidly diversified when placed in the context of the repeated environmental fluctuations that occurred throughout the mid and late Miocene (Michel et al. 2014, Hunt 2016) before becoming functionally canalized through development and stabilizing selection. This pattern is consistent with a scenario in which morphological features evolve by stochastic diffusion on a Simpsonian adaptive landscape that is alternately dampened and released by relaxed and shifting patterns in character integration.

Rapid and frequent environmental fluctuations throughout the Miocene suggest a possible abundance of available adaptive zones during the evolution of both cercopithecoids and hominoids. Paleoecological information extracted from deposits containing key hominoid taxa are so variable that environmental turnover often outpaces the effects of time-averaging (Michel et al. 2014). Early Miocene ape and monkey species evolved in overlapping environmental conditions and shared similar ecomorphological and dietary habits. However, as their lineages diverged, apes evolved a diverse set of locomotor suites, while evolution in locomotor features has remained comparatively static across old world monkeys (Hunt 2016). The patterns in developmental and mosaic morphological evolution revealed here suggest that the substantial differences between the two taxa in phenotypic disparity were shaped more by differences in patterns in integration and constraint rather than from the ecological opportunities available to either. While hominoids and cercopithecoids inhabited a similar range of habitats throughout their evolution, and so likely experienced a similar abundance of ecological opportunity, greater constructional opportunity is a distinguishing feature that correlates with hominoids’ vast disparity in skeletal morphology.

The scenario entertained here for apes is consistent with recent work that suggests that the initial burst in phenotypic disparity that often accompanies the origin of a new lineage can result from an early relaxation of constraints through the dissolution of integrated modules that is followed by reformation of character suites that lead to rapid, correlated evolutionary changes between constituent characters (Wagner 2018). In the cited work, the author distinguishes this ‘breakup-relinkage’ model with one where evolutionary rates are elevated by a relaxation of constraint alone. The ‘breakup-relinkage’ model is directly supported by the phylogenetic integration pattern, which show that an initial decrease in integration at the root of hominoids was followed by an increase at the following node (Fig. 2). Therefore, the recanalization of functional variation into a set of suites with distinct functional and developmental properties early in hominoid variation is likely equally important to the initial relaxation in constraint in having driven the remarkable morphological disparity observed across hominoid species. *“Preadaptation” and developmental enablers in apes*

Bursts in phenotypic change and diversification are often preceded by evolutionary enablers that facilitate the construction of more diverse morphological forms or fulfill a necessary condition for the later emergence of a more derived trait. Such patterns have been suggested to stem from higher-level processes variously referred to as “preadaptation” (Bock 1959), “exaptation” (Gould and Vrba 1982), and “developmental enablement” (Donoghue 2005). The evolution of developmental, morphological, and behavioral traits is often hierarchical through functional inter-dependencies and temporally autocorrelated, with changes in traits at relatively lower levels ‘setting the stage’ for higher-level traits by providing the developmental or structural conditions necessary for their emergence. The stepwise macroevolutionary bursts observed here show that hominoid evolution has been defined by sequential, episodic releases in constraint that likely set the stage for later innovations that compounded upon earlier rearrangements in the structure of developmental and functional integration.

The related concepts of preadaptation, exaptation, and developmental enablement all focus on identifying the origin of single characters that facilitate the later emergence of more derived states. However, the patterns presented here suggest that such processes may often be better characterized as reflecting larger changes in developmental and functional character-linkage systems rather than by the sequential derivation of individual characters. The dramatic bursts in evolutionary rate that occur in the largest character suites coincide with or immediately follow the reduction in integration that occurred at the root of hominoids. This suggests that a general shift in functional and/or developmental integration occurred near this time. This shift precedes and, although speculative, may have resulted in a general predisposition across apes that facilitated the later functional divergence of later species into an unusually broad range of habits. Therefore, rather than stemming from a single “preadaptive” character, the morphological disparity of apes and the emergence of features derived in single lineages (such as bipedalism in humans) may be better explained by a shift in developmental and functional integration patterns earlier in ape evolution.

Enhanced evolvability conferred by shifts in the structure of integration contributed to the emergence of key human traits. The apparent formation of suites C2 and C4, represented disproportionately by the cranium and pelvis, along the branch leading to humans and chimpanzees coincided with an initial burst in evolutionary rate. Both of these events immediately preceded the dramatic bursts in *Homo* (Fig. 3). Likewise, EQ and ED have been evolving rapidly throughout the ape radiation, with ED experiencing an initial increase in evolutionary rate before the splitting of gorillas, chimps, and humans followed by more dramatic upticks of rate in humans in both traits (Fig. 5c). While distinct in its details, this pattern in chimps and humans provides further evidence for the preadaptive potency of shifts in systems of modularity and integration in driving dramatic anatomical rearrangements. It also illustrates that the broad range of possible evolutionary pathways opened by either an overall reduction in integration (at the hominoid root) or the formation of new modules (in chimps and humans), rather than movement toward a specific set of phenotypes, is the means through which such predispositions facilitate evolutionary remodeling.

It is possible that the initial bursts in ED and suite C2 and C3 evolution, coupled with the reduced integration in the MRCA of chimp and humans, may reflect a set of structural rearrangements that facilitated, at least in part, the later emergence of human-specific neurological and cognitive features. This scenario echoes the results of Rice (1998), who demonstrated that humans and chimpanzees, to the exclusion of gorillas, both inherited a shared ‘additional’ stage of brain development from our most recent common ancestor. It is therefore possible that, in addition to sharing a similar derived structure of modularity, chimpazees possess more of the neurological and cranial groundwork that contributed to the evolution of humans’ particular mode of cognition than has previously been appreciated.

The patterns uncovered here suggest that the concepts of preadapation and exaptation may sometimes be more compellingly applied by shifting focus from individual features to patterns in integration and modularity across broad suites of characters. Although the concept is essentially unchanged as applied to integration patterns, it may be more accurate in the example here to refer instead to an ‘evolutionary predisposition’ given the lack of evidence for a specific sequence of selective regimes (Smith et al. 2018). This semantic argument is likely also valid in more traditional examples of single trait exaptations or preadaptations. And so, a generalization from preadaptation or exaptation to evolutionary predisposition that 1) incorporates integration patterns and 2) shifts emphasis from adaptation, which makes assumptions about the (often unknowable) set of historical microevolutionary processes, to the sequence of observable states in both characters and integration patterns can provide a stronger conceptual basis for future work that considers contingency and sequential order in the evolutionary pathways of complex traits. Although this reconceptualization might superficially appear to de-emphasize evolutionary process in favor of pattern, I would instead argue that constraining the scale of inquiry and incorporating a broader range of biological complexity can facilitate the reconstruction of elusive higher-level macroevolutionary processes that shape organismal diversification and disparification over deep timescales. This formulation also avoids the conceptual and practical frustrations that occur when attempting to invoke the concepts of preadaptation and exaptation to explain evolutionary history over deep timescales by reducing dependency on knowledge of lower-level population processes.

### Diversification and disparification in haplorrhine evolution

Cercopithecoids have diversified in species number at over twice the rate as hominoids over the same time scale (Table 1). One limitation of these analyses stems from cercopithecoids’ somewhat poor fossil record as compared to hominoids. Nevertheless, the difference in preservation rates estimated in each clade (∼3 in apes compared to ∼2.6 in cercopithecoids) should ameliorate some of the potential effect of this bias on parameter estimates. The general pattern of higher net diversification in cercopithecoids is also consistent with previous neontological results (Purvis et al. 1995), suggesting that the paleontological estimates presented here are adequately robust to these effects. This difference in lineage diversification rates between both taxa demonstrates that increased morphological disparification is, overall, not correlated with faster lineage diversification in haplorrhine primates. While the evolutionary predisposition conferred upon apes by their relaxed integration corresponds to an increase in morphological disparity, this effect does not extend to lineage diversity.

On the surface, the overall similarity in the environments, dietary niches, and geographical ranges of hominoids and cercopithecoids suggests that cercopithecoids have a selective advantage in the species-sorting dynamics between the two clades. However, the long-term persistence of hominoids and their remarkable innovation in locomotor function relative to cercopithecoids throughout this timespan suggests an equilibrium in their relative abundances (Van Valen 1975, Chesson 2000). Although cercopithecoids experienced less dramatic innovation in postcranial morphology, they display a broader range of derived digestive physiologies, including both intestinal and dental features associated with food processing (Hunt 2016). Given these observations, it is possible that the differential in diversification rate between cercopithecoids and hominoids stems from differences in the timescales of their life histories and other factors not directly related to interspecific competitive ability. If this is the case, hominoids may achieve steady persistence through higher evolvability conferred by reduced integration and increased developmental flexibility when faced with changing environments, while cercopithecoids may do so through a combination of their higher net diversification rates and capability to exploit a range of dietary resources through their digestive physiologies. Such differences would be consistent with an equilibrium model of coexistence rather than a simple species selection model that predicts the ultimate extinction of hominoids through competitive exclusion by cercopithecoids. Nevertheless, more rigorous comparison of each of these attributes in both lineages is needed to further constrain the range of possible explanatory factors.

### Constrained Markovian diffusion and entropy

The patterns in character disparity shown here evoke a Markovian diffusion that is dampened by constraint. The tightly integrated nature of vertebrate body plans suggests major constraints in their ability to fill morphospace. However, the pattern displayed by apes suggests that the structure of integration may sometimes shift, allowing diffusion into new areas. Morphological evolution might then be conceived as a multivariate diffusion in morphospace where movement is directionally biased due to fitness differences modelled by a traditional Simpsonian adaptive landscape and certain regions are rendered inaccessible by structural interactions between characters. Although this diffusive model, including the feedback links between phenotype, development, and environment, has been considered previously (Fisher 1986), its behavior has not been well explored in this context.

The Monte Carlo simulations presented here provide a theoretical illustration of the population-level dynamics stemming from one possible set of integration scenarios and their potential effects on the evolutionary disparity/rate along a single branch in Figures 3 or 4. The results shed light on the empirical analyses by demonstrating the overlapping and distinct empirical patterns that can result from varying levels of developmental and functional integration in a simple closed system in the absence of environmental fluctuation (Fig. 6). As is expected from quantitative genetic theory (Cheverud 1984, Maynard-Smith et al. 1985), developmental and functional constraints can generate similar patterns (Fig. 6a and 6b). In all these cases, the overall entropy observed in the system over time remains low, with the system tending to become stuck or moving away very slowly from a suboptimal location (Fig 4c and 4e) or crawling slowly in concert along a gradient (Fig 6a and 6b). However, when both types of constraint are released, the system displays high entropy, with each trait able to independently jump between peaks (Fig. 6d). Such unconstrained movement would be expected to result in higher phenotypic disparity over macroevolutionary timescales by freeing individual lineages to generate novel character combinations when exposed to distinct and changing environments.

**Figure 6.**
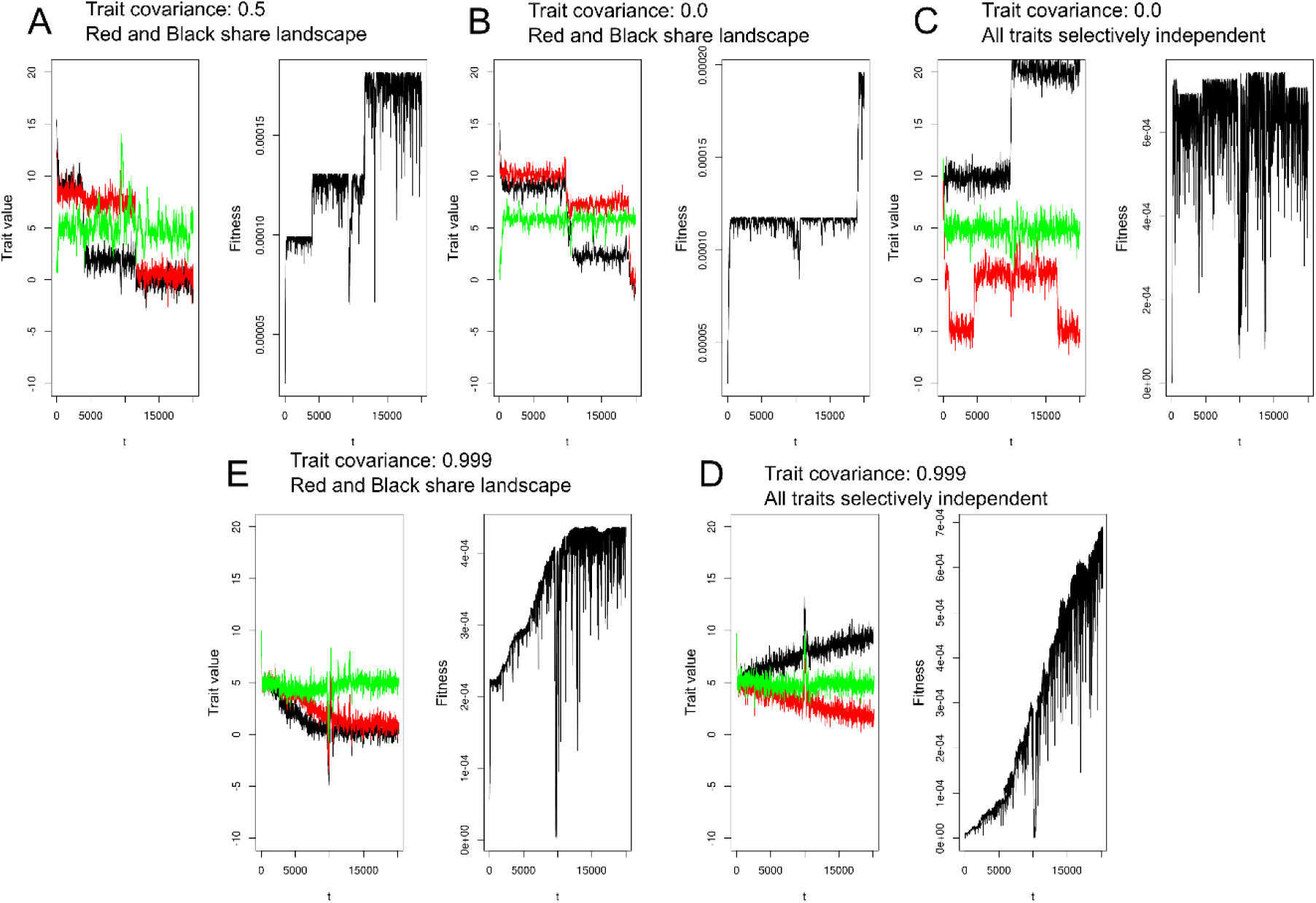
Monte Carlo simulations of integrated and atomized systems of three continuous traits. Each box is a simulated diffusion between three traits under A) moderate developmental linkage and shared selection, B) no developmental linkage and shared selection, C) no developmental linkage and selective independence, D) high developmental linkage and selective independence, and E) high developmental linkage and shared selection.

The patterns in increased evolutionary rate and entropy across traits displayed by the less constrained simulated diffusions provide mechanistic explanations that are consistent with the bursts in evolutionary rate encountered throughout hominoid evolution. Although it is not possible in the empirical example to distinguish between functional and developmental integration as is done in the simulations, the general pattern of reduced integration that gives way to increased evolutionary rate is consistent between the two. Alternation between the varying strengths of integration as explored in the simulations would likely generate substantial novelty during periods of relaxed integration. Episodes of reduced developmental constraint (Figs. 2 and 5a) may have generated an initial burst of constructional opportunity that lead to rapid morphological diversification as ape populations were exposed to highly variable environments throughout the mid-Miocene. If the processes modelled in the simulations drove the patterns revealed in the empirical analyses, this sequence of events would represent true preadaptation in the sense that relaxed integration leads to increased mean population fitness. Under the evolutionary models explored here and elsewhere (Wagner 2018), stabilizing effects such as developmental canalization (Waddington 1959) or functional covariance (Cheverud 1984) would then be expected to re-constrain the newly divergent phenotypes after an initial burst in disparification. This dampened diffusive model was explored in early work in theoretical morphology that focused on the constrained filling of morphospace (Raup 1968).

The simulated diffusions and empirical pattern in evolutionary predisposition revealed here hint at the source of teleological concepts in human evolution. If the morphological and neurological traits can be assumed to have followed a multivariate Markovian diffusion that is at least qualitatively similar to the simulations in Figure 6 (especially Fig. 6a), the pattern in evolutionary rates along the path leading to humans may paint a misleading view that such changes have followed a progressive trend leading to a human-defined apex. However, the mosaic analysis here suggests that humans’ biological uniqueness can be attributed to a relatively small number of anatomical rearrangements built over a longer diffusion into an area of developmental-morphospace occupied by all apes that is characterized by increased structural opportunity. Instead of treating humans as exceptional, this view suggests that the structural opportunities that emerged early in ape evolution may have freed the ancestors of currently extant taxa to blaze unique trajectories along a complex, multivariate adaptive landscape that would have been otherwise inaccessible. Structurally, all ape species might therefore be viewed as equally progressive in their evolution, with species-level differences in form and function shaped by stochastic differences in separate realized evolutionary paths along a shared adaptive landscape. Alternatively, separate taxa may have forged new and unique adaptive landscapes following speciation events as structural differences between newly isolated species drove the creation of new functional niches early in hominoid evolution. The derived structure of integration and neurological development shared by chimps and humans highlights the importance of contingency in human evolution—chimps’ sharing of much of the same underlying constructional architecture shows how this derived predisposition allowed, but did not determine, the emergence of humans’ unique suite of phenotypes.

The empirical and simulated analyses presented here evoke a picture of morphological evolution that involves episodic diffusion across adaptive zones. However, the empirical and theoretical analyses allow for alternating and interacting roles for both ecological and constructional opportunity. This view can supplement existing conceptions of adaptive radiation and key innovation by providing an alternative to ecological opportunity as a lone driving factor in the disparification of form. Instead, phenotypic disparity produced through diffusion into a set of abundant adaptive zones can be directed and limited by the availability of structural opportunity. Nature surely merges these simplified extremes when shaping patterns in biodiversity (Seilacher 1991), and so further integrative study of the interplay between ecological and constructional factors will be critical in developing a pluralistic understanding of the complex patterns and processes that have shaped the vast diversity across the tree of life.

## Acknowledgements

I wish to thank D Alvarado-Serrano, M Cosman, CW Dick, DC Fisher, M Foote, E Greiner, D Jablonski, LM MacLatchy, H Marx, J Saulsbury, SA Smith, G Slater, N Walker-Hale, M Webster for conversations that have greatly benefitted this work. I also thank S Worthington for help accessing his skeletal measurements that comprised a large portion of the character matrix analyzed here. The manuscript benefitted immensely from careful reviews provided by P Wagner and PD Polly.

---

**Table S1:**
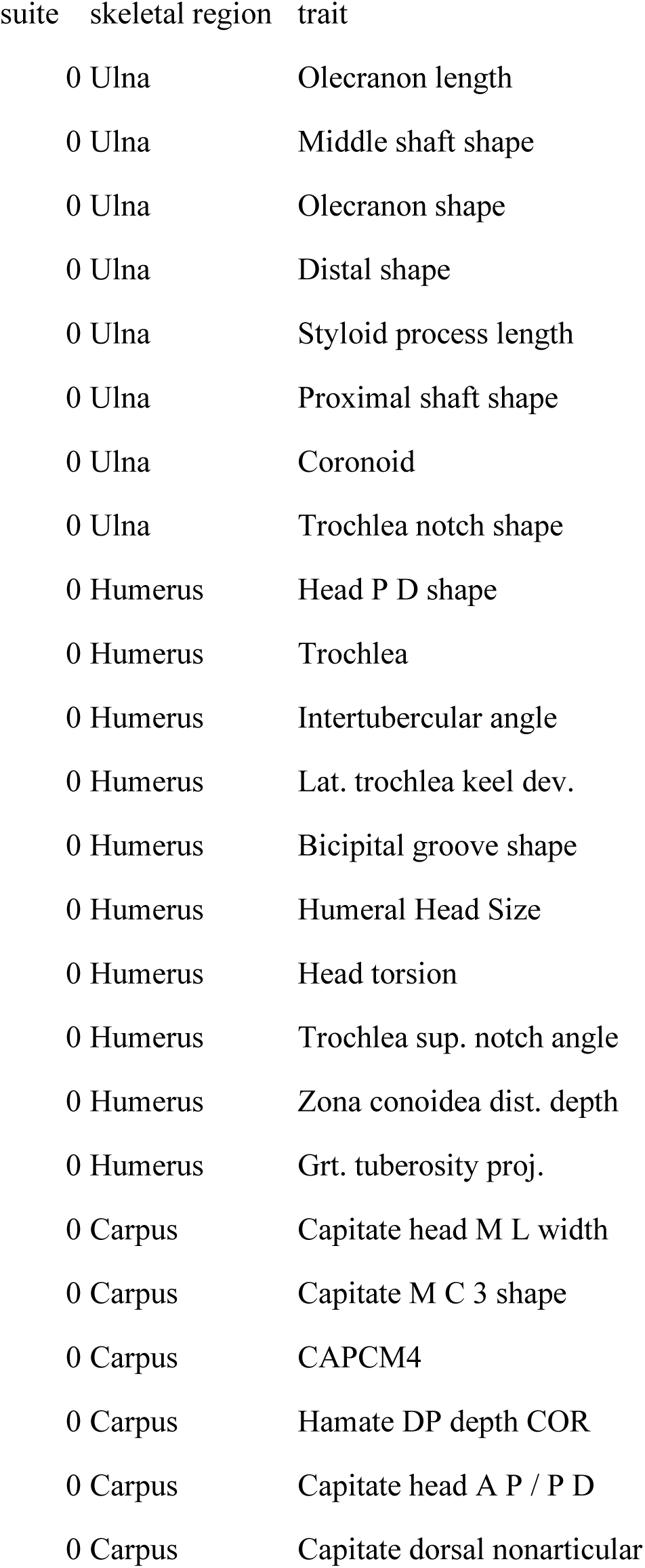
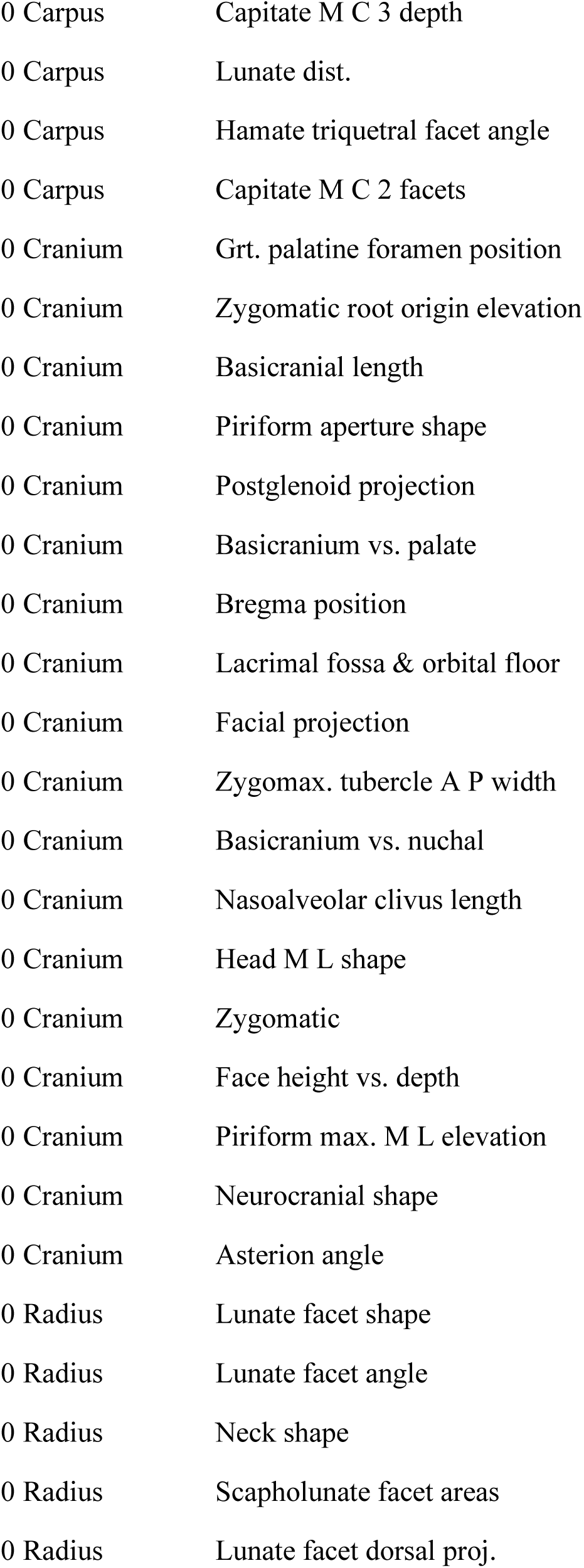
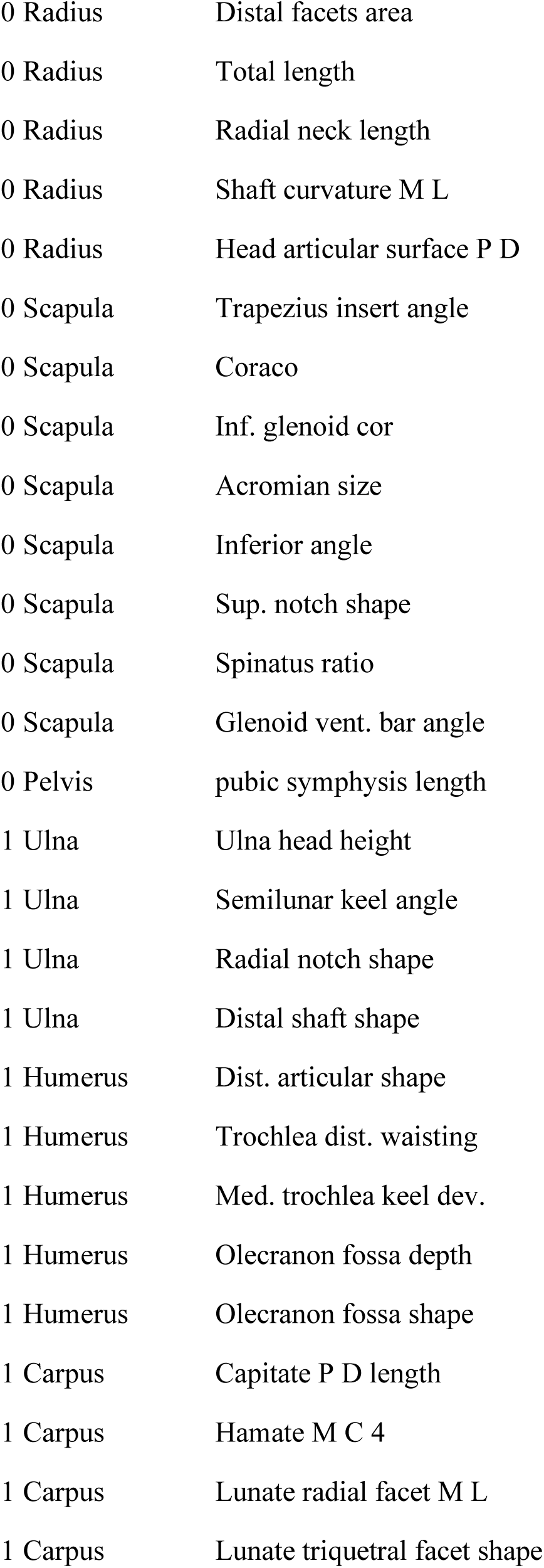
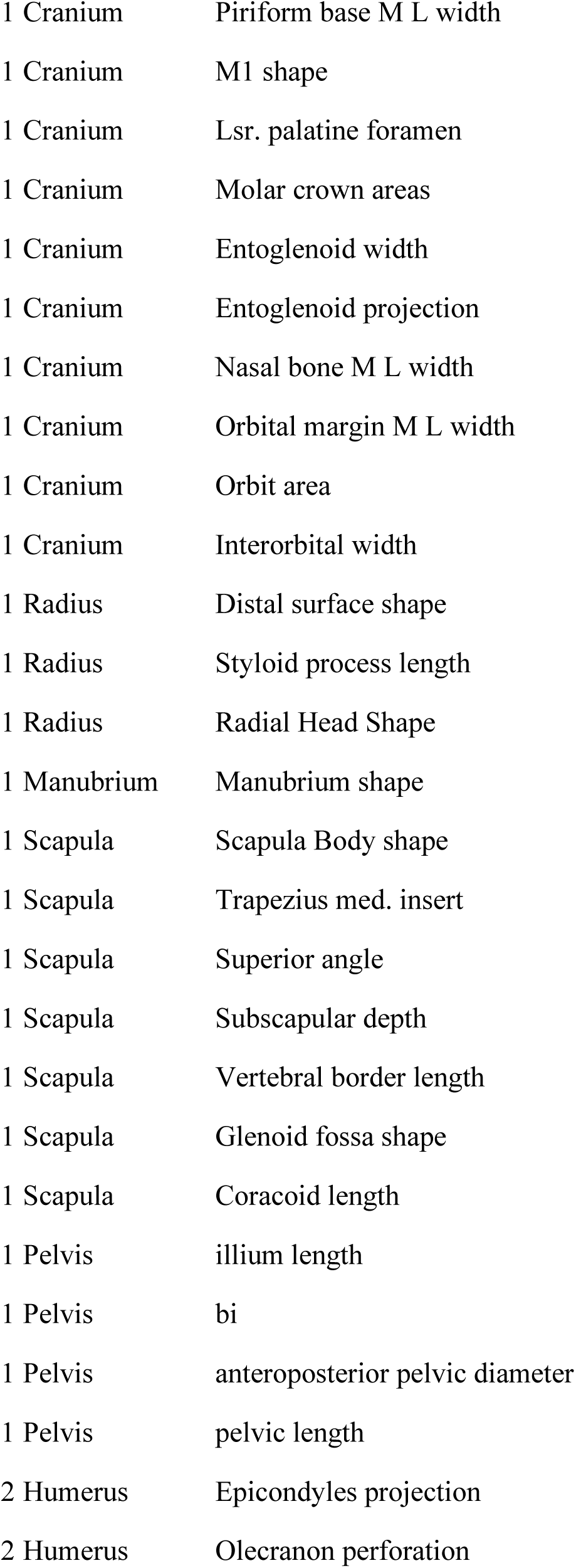
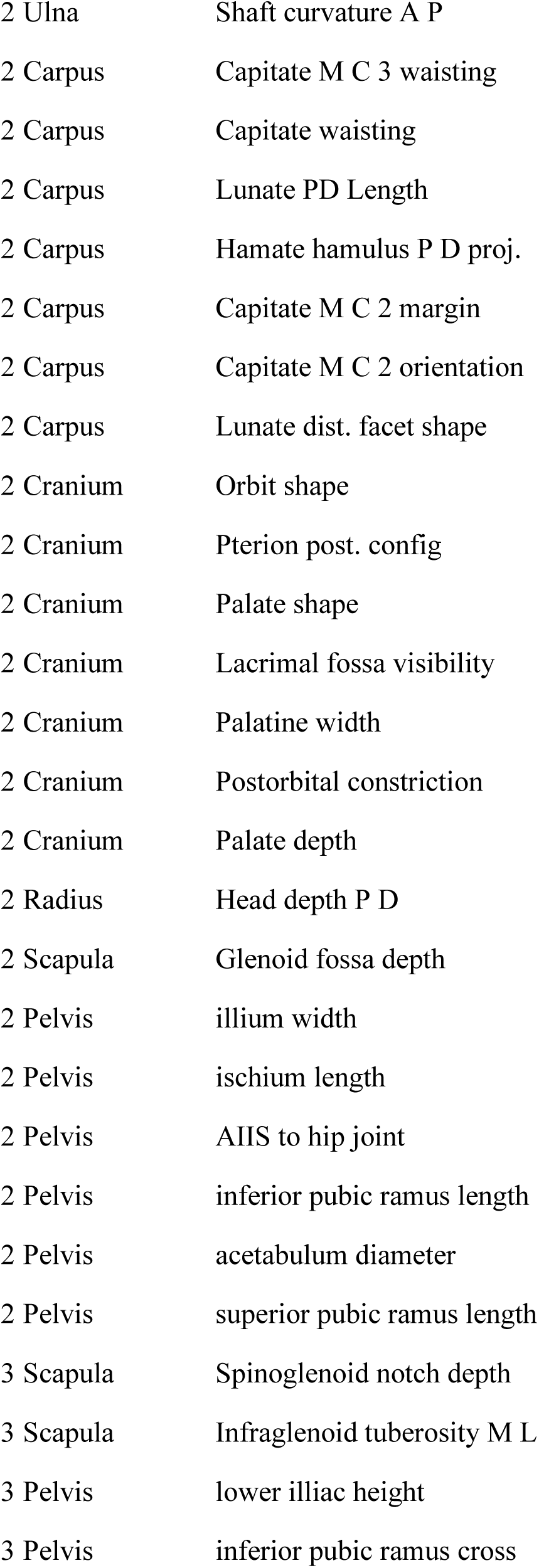
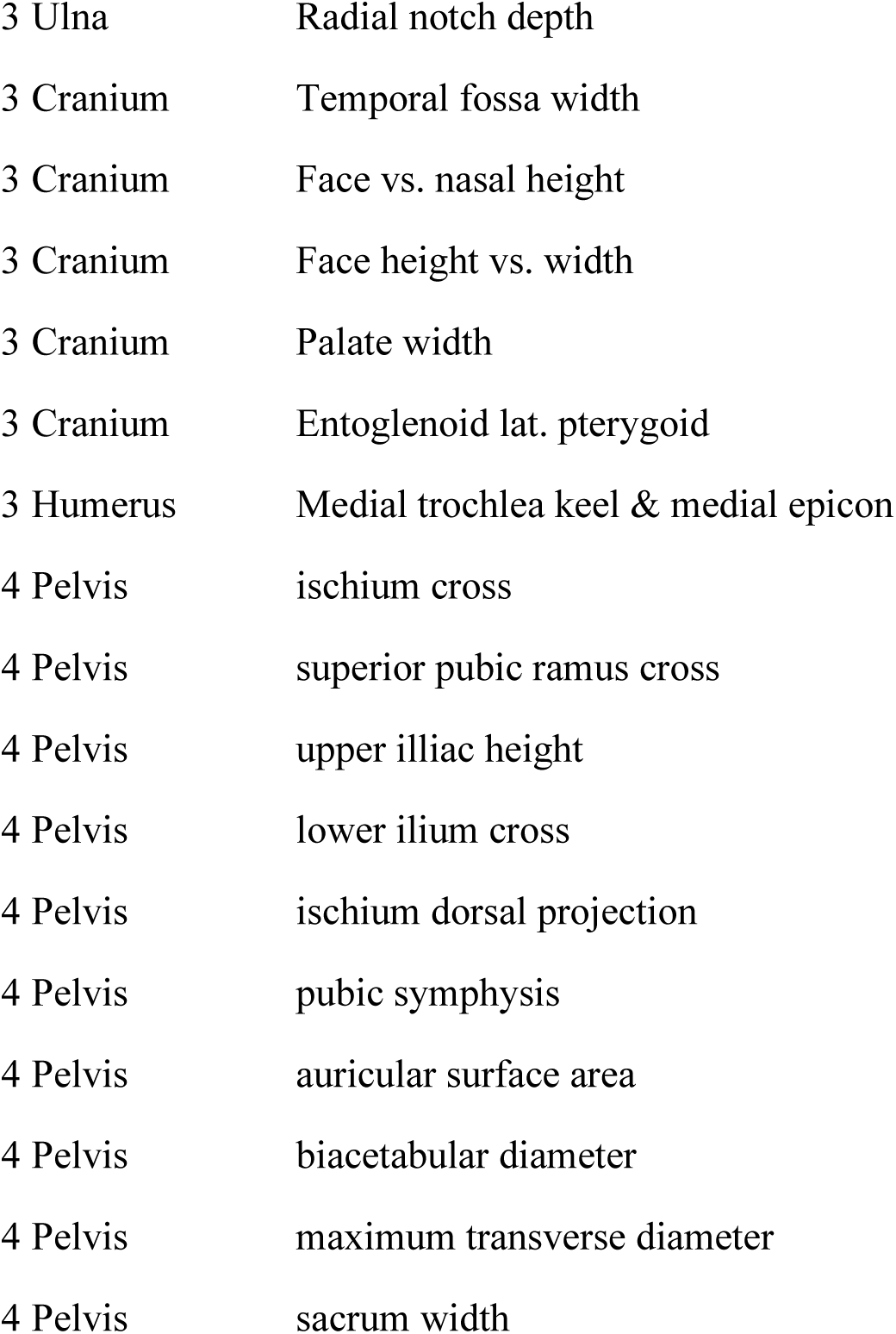
List of traits and their skeletal regions that comprise each of the mosaic suites detected from the skeletal data.

| suite | skeletal region | trait |
| --- | --- | --- |
| 0 | Ulna | Olecranon length |
| 0 | Ulna | Middle shaft shape |
| 0 | Ulna | Olecranon shape |
| 0 | Ulna | Distal shape |
| 0 | Ulna | Styloid process length |
| 0 | Ulna | Proximal shaft shape |
| 0 | Ulna | Coronoid |
| 0 | Ulna | Trochlea notch shape |
| 0 | Humerus | Head P D shape |
| 0 | Humerus | Trochlea |
| 0 | Humerus | Intertubercular angle |
| 0 | Humerus | Lat. trochlea keel dev. |
| 0 | Humerus | Bicipital groove shape |
| 0 | Humerus | Humeral Head Size |
| 0 | Humerus | Head torsion |
| 0 | Humerus | Trochlea sup. notch angle |
| 0 | Humerus | Zona conoidea dist. depth |
| 0 | Humerus | Grt. tuberosity proj. |
| 0 | Carpus | Capitate head M L width |
| 0 | Carpus | Capitate M C 3 shape |
| 0 | Carpus | CAPCM4 |
| 0 | Carpus | Hamate DP depth COR |
| 0 | Carpus | Capitate head A P / P D |
| 0 | Carpus | Capitate dorsal nonarticular |

**Table S1:** List of traits and their skeletal regions that comprise each of the mosaic suites detected 875 from the skeletal data.
|  |  |
| --- | --- |
| 0 Carpus | Capitate M C 3 depth |
| 0 Carpus | Lunate dist. |
| 0 Carpus | Hamate triquetral facet angle |
| 0 Carpus | Capitate M C 2 facets |
| 0 Cranium | Grt. palatine foramen position |
| 0 Cranium | Zygomatic root origin elevation |
| 0 Cranium | Basicranial length |
| 0 Cranium | Piriform aperture shape |
| 0 Cranium | Postglenoid projection |
| 0 Cranium | Basicranium vs. palate |
| 0 Cranium | Bregma position |
| 0 Cranium | Lacrima fossa & orbital floor |
| 0 Cranium | Facial projection |
| 0 Cranium | Zygomax. tubercle A P width |
| 0 Cranium | Basicranium vs. nuchal |
| 0 Cranium | Nasoalveolar clivus length |
| 0 Cranium | Head M L shape |
| 0 Cranium | Zygomatic |
| 0 Cranium | Face height vs. depth |
| 0 Cranium | Piriform max. M L elevation |
| 0 Cranium | Neurocranial shape |
| 0 Cranium | Asterion angle |
| 0 Radius | Lunate facet shape |
| 0 Radius | Lunate facet angle |
| 0 Radius | Neck shape |
| 0 Radius | Scapholunate facet areas |
| 0 Radius | Lunate facet dorsal proj. |
| 0 Radius | Distal facets area |
| 0 Radius | Total length |
| 0 Radius | Radial neck length |
| 0 Radius | Shaft curvature M L |
| 0 Radius | Head articular surface P D |
| 0 Scapula | Trapezius insert angle |
| 0 Scapula | Coraco |
| 0 Scapula | Inf. glenoid cor |
| 0 Scapula | Acromian size |
| 0 Scapula | Inferior angle |
| 0 Scapula | Sup. notch shape |
| 0 Scapula | Spinatus ratio |
| 0 Scapula | Glenoid vent. bar angle |
| 0 Pelvis | pubic symphysis length |
| 1 Ulna | Ulna head height |
| 1 Ulna | Semilunar keel angle |
| 1 Ulna | Radial notch shape |
| 1 Ulna | Distal shaft shape |
| 1 Humerus | Dist. articular shape |
| 1 Humerus | Trochlea dist. waisting |
| 1 Humerus | Med. trochlea keel dev. |
| 1 Humerus | Olecranon fossa depth |
| 1 Humerus | Olecranon fossa shape |
| 1 Carpus | Capitate P D length |
| 1 Carpus | Hamate M C 4 |
| 1 Carpus | Lunate radial facet M L |
| 1 Carpus | Lunate triquetral facet shape |
| 1 Cranium | Piriform base M L width |
| 1 Cranium | M1 shape |
| 1 Cranium | Lsr. palatine foramen |
| 1 Cranium | Molar crown areas |
| 1 Cranium | Entoglenoid width |
| 1 Cranium | Entoglenoid projection |
| 1 Cranium | Nasal bone M L width |
| 1 Cranium | Orbital margin M L width |
| 1 Cranium | Orbit area |
| 1 Cranium | Interorbital width |
| 1 Radius | Distal surface shape |
| 1 Radius | Styloid process length |
| 1 Radius | Radial Head Shape |
| 1 Manubrium | Manubrium shape |
| 1 Scapula | Scapula Body shape |
| 1 Scapula | Trapezius med. insert |
| 1 Scapula | Superior angle |
| 1 Scapula | Subscapular depth |
| 1 Scapula | Vertebral border length |
| 1 Scapula | Glenoid fossa shape |
| 1 Scapula | Coracoid length |
| 1 Pelvis | illium length |
| 1 Pelvis | bi |
| 1 Pelvis | anteroposterior pelvic diameter |
| 1 Pelvis | pelvic length |
| 2 Humerus | Epicondyles projection |
| 2 Humerus | Olecranon perforation |
| 2 Ulna | Shaft curvature A P |
| 2 Carpus | Capitate M C 3 waisting |
| 2 Carpus | Capitate waisting |
| 2 Carpus | Lunate PD Length |
| 2 Carpus | Hamate hamulus P D proj. |
| 2 Carpus | Capitate M C 2 margin |
| 2 Carpus | Capitate M C 2 orientation |
| 2 Carpus | Lunate dist. facet shape |
| 2 Cranium | Orbit shape |
| 2 Cranium | Pterion post. config |
| 2 Cranium | Palate shape |
| 2 Cranium | Lacrima fossa visibility |
| 2 Cranium | Palatine width |
| 2 Cranium | Postorbital constriction |
| 2 Cranium | Palate depth |
| 2 Radius | Head depth P D |
| 2 Scapula | Glenoid fossa depth |
| 2 Pelvis | ilium width |
| 2 Pelvis | ischium length |
| 2 Pelvis | AIIS to hip joint |
| 2 Pelvis | inferior pubic ramus length |
| 2 Pelvis | acetabulum diameter |
| 2 Pelvis | superior pubic ramus length |
| 3 Scapula | Spinoglenoid notch depth |
| 3 Scapula | Infraglenoid tuberosity M L |
| 3 Pelvis | lower iliac height |
| 3 Pelvis | inferior pubic ramus cross |
| 3 Ulna | Radial notch depth |
| 3 Cranium | Temporal fossa width |
| 3 Cranium | Face vs. nasal height |
| 3 Cranium | Face height vs. width |
| 3 Cranium | Palate width |
| 3 Cranium | Entoglenoid lat. pterygoid |
| 3 Humerus | Medial trochlea keel & medial epicon |
| 4 Pelvis | ischium cross |
| 4 Pelvis | superior pubic ramus cross |
| 4 Pelvis | upper illiac height |
| 4 Pelvis | lower ilium cross |
| 4 Pelvis | ischium dorsal projection |
| 4 Pelvis | pubic symphysis |
| 4 Pelvis | auricular surface area |
| 4 Pelvis | biacetabular diameter |
| 4 Pelvis | maximum transverse diameter |
| 4 Pelvis | sacrum width |

## Notes

#### Summary of Updates

Major edits to text

